# Nanomechanical phenotypes in cMyBP-C mutants that cause hypertrophic cardiomyopathy

**DOI:** 10.1101/2020.09.19.304618

**Authors:** Carmen Suay-Corredera, Maria Rosaria Pricolo, Diana Velázquez-Carreras, Carolina Pimenta-Lopes, David Sánchez-Ortiz, Iñigo Urrutia-Irazabal, Silvia Vilches, Fernando Dominguez, Giulia Frisso, Lorenzo Monserrat, Pablo García-Pavía, Elías Herrero-Galán, Jorge Alegre-Cebollada

**Author notes:** To whom correspondence should be addressed. Twitter: @AlegreCebollada.

## Abstract

Hypertrophic cardiomyopathy (HCM) is a disease of the myocardium caused by mutations in sarcomeric proteins with mechanical roles, such as the molecular motor myosin. Around half of the HCM-causing genetic variants target contraction modulator cardiac myosin-binding protein C (cMyBP-C), although the underlying pathogenic mechanisms remain unclear since many of these mutations cause no alterations in protein structure and stability. As an alternative pathomechanism, here we have examined whether pathogenic mutations perturb the nanomechanics of cMyBP-C, which would compromise its modulatory mechanical tethers across sliding actomyosin filaments. Using single-molecule atomic force spectroscopy, we have quantified mechanical folding and unfolding transitions in cMyBP-C mutant domains. Our results show that domains containing mutation R495W are mechanically weaker than wild-type at forces below 40 pN, and that R502Q mutant domains fold faster than wild-type. None of these alterations are found in control, non-pathogenic variants, suggesting that nanomechanical phenotypes induced by pathogenic cMyBP-C mutations contribute to HCM development. We propose that mutation-induced nanomechanical alterations may be common in mechanical proteins involved in human pathologies.

## INTRODUCTION

Hypertrophic cardiomyopathy (HCM) is the most common inherited cardiac muscle disease, affecting up to 1 in 200 individuals ^2-3^. Macroscopically, HCM is characterized by thickened left ventricular walls and reduced size of the left ventricular chamber, while at the tissue level, HCM myocardium typically shows interstitial fibrosis and fiber disarray (**Figure 1a**). These structural changes occur alongside functional defects such as diastolic dysfunction, which can lead to the most severe consequences of the disease including heart failure and sudden cardiac death ^4-6^. Despite encouraging advances ^7^, currently there are no therapies to revert nor prevent HCM pathogenesis and clinical management relies on long-term palliative treatments and surgical procedures ^4-5^.

**Figure 1.**
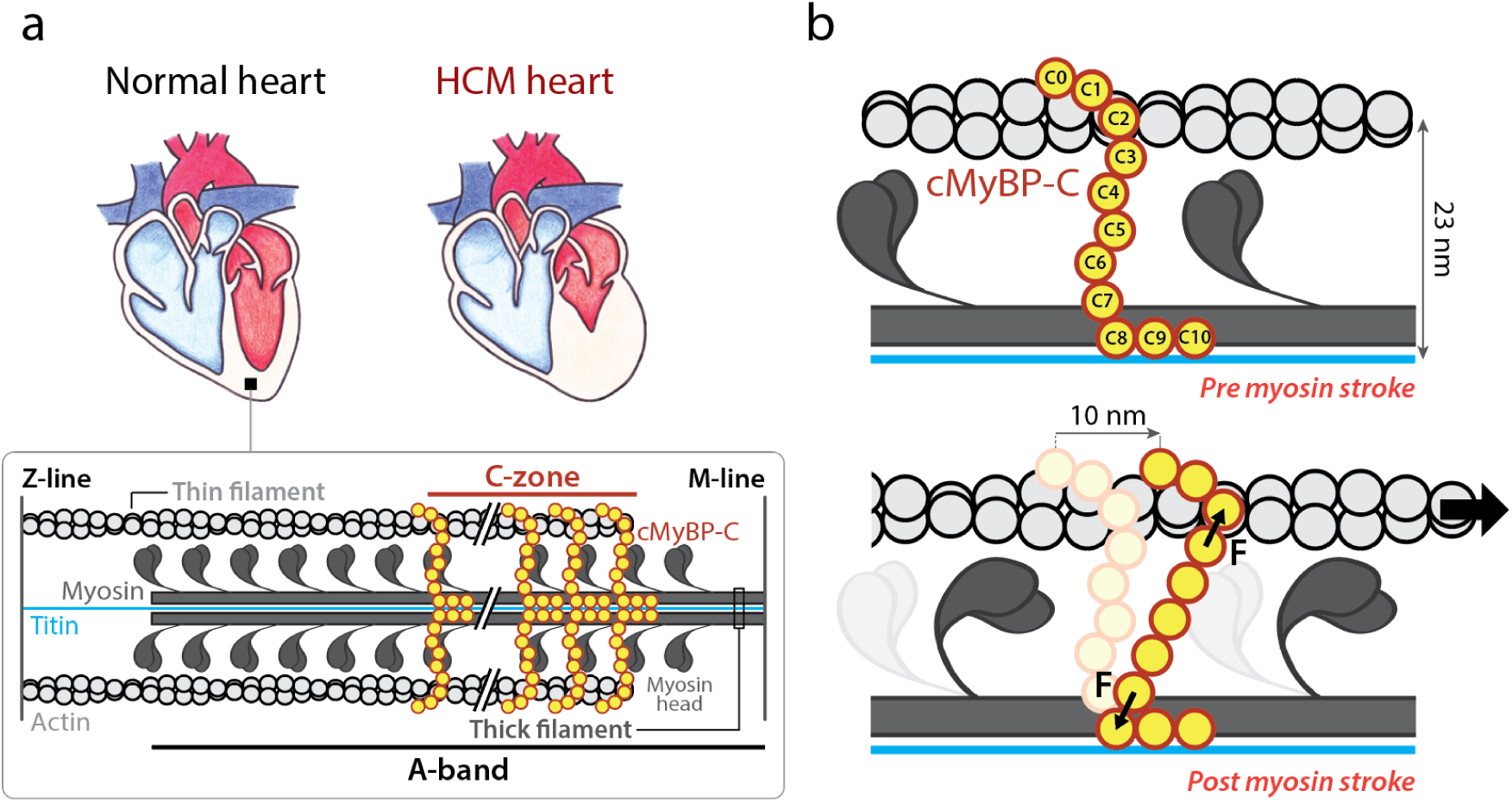
Overview of the mechanical role of cMyBP-C in the sarcomere. **(a)** Comparison of a healthy heart and an HCM counterpart, which shows thicker left ventricular walls and reduced left ventricle volume. *Inset*: schematics of the sarcomere, whose contraction relies on actin-based thin filaments that glide over myosin-containing thick filaments thanks to myosin power strokes. cMyBP-C (in yellow) is located in the C-zone, a part of the A-band of the sarcomere. The M-line and the Z-line structures, which arrange filaments supporting sarcomere organization, are also shown ^1^. **(b)** cMyBP-C tethers are subject to mechanical force during a 10 nm myosin power stroke. Interfilament distance is indicated.

The majority of HCM cases are caused by autosomal dominant mutations targeting mechanical proteins of the sarcomere, the basic contractile unit of cardiomyocytes ^5, 8-9^ (**Figure 1a**). In sarcomeres, myosin heads use the energy coming from ATP hydrolysis to extend from the myosin backbone in the thick filaments, establish cross-bridges with the neighboring actin thin filaments and generate ∼10-nm power strokes that propel the thin filaments past the thick ones leading to muscle contraction (**Figure 1b**) ^4, 8, 10-12^. Cardiac myosin-binding protein C (cMyBP-C) is a well-known negative modulator of sarcomere contraction by complex mechanisms that are not fully understood ^13-15^. This multidomain protein is located in the C-zone of the sarcomere, grouped in nine regularly-spaced transverse stripes that are 43 nm apart from each other ^16^. cMyBP-C’s C-terminal C8-C10 domains run axially along the thick filament, interacting both with the myosin backbone and titin. The central and N-terminal domains of the protein extend radially from the thick filament towards the thin filament located ∼23-nm away (**Figure 1**) ^13, 15-20^. In this geometry, the N-terminal domains of cMyBP-C can interact both with the myosin head region and with the actin filament. Since the lifetime of the interaction between cMyBP-C and actin filaments is on the order of 20 −300 ms ^21^ and the unloaded velocity of contraction per half sarcomere can be as high as 7 nm/ms ^22^, myosin power strokes are expected to induce substantial mechanical strain on cMyBP-C, which therefore contributes a viscous load that opposes actomyosin contraction ^23^ (**Figure 1b**). Hence, how cMyBP-C tethers respond to mechanical load ^24-26^ can be fundamental for its modulatory role on sarcomere power generation.

*MYH7* and *MYBPC3*, which codify for β-myosin heavy chain and cMyBP-C, respectively, are the most frequently mutated genes in HCM, accounting for 80% of cases ^13, 15, 19^. A large proportion of pathogenic variants in *MYBPC3* result in cMyBP-C truncations that induce HCM via reduction in total cMyBP-C content (protein haploinsufficiency) ^4, 13, 28-34^. However, many *MYBPC3* pathogenic mutations only cause single amino acid substitutions that result in full-length, mutant cMyBP-C proteins that can incorporate into sarcomeres to the same levels as the wild-type protein ^19, 29, 31^. The molecular deficits of these missense mutations remain largely unexplored. Some of them have been proposed to disrupt cMyBP-C interaction with actomyosin filaments ^35-37^ or to induce extensive protein destabilization ^38-39^; however, many HCM-causing missense mutants appear to operate via alternative, unidentified mechanisms ^40^. Prompted by the ability of cMyBP-C to establish mechanical tethers that modulate sarcomere contraction (**Figure 1b**), we hypothesized that HCM-causing mutations may perturb the nanomechanics of cMyBP-C leading to altered sarcomere activity. Here, we have used single-molecule force spectroscopy by atomic force microscopy (AFM) to test this hypothesis ^41^. We have found that HCM-causing mutations can affect the mechanical stability and folding rate of the targeted domains, raising the possibility that alteration of cMyBP-C nanomechanics contributes to HCM pathogenesis.

## RESULTS

### Selection of pathogenic and non-pathogenic cMyBP-C variants

Following strict assignment of pathogenicity based on clinical and epidemiological data ^40^, we selected 4 pathogenic missense mutations (R495Q, R495W, R502Q and R502W) and 2 non-pathogenic variants (G507R and A522T) targeting *MYBPC3*. The variants are located in exon 17 of the gene, which together with exon 16 codifies for the C3 domain of cMyBP-C (**Supplementary Text S1**; **Figure 2**) ^42^. No protein interactors have been described for this central domain, as expected from the location of C3 far from cMyBP-C’s anchoring points to actomyosin filaments ^43-44^. Hence, mutations targeting C3 are arguably not predicted to affect cMyBP-C interactions, which is in agreement with the normal sarcomere localization of cMyBP-C missense variants in HCM myocardium ^31^. We first verified that the pathogenic mutations do not induce defects in RNA splicing or extensive protein structural destabilization, two classical protein haploinsufficiency drivers linked to pathogenicity in 45% of cMyBP-C missense mutations ^40^.

**Figure 2.**
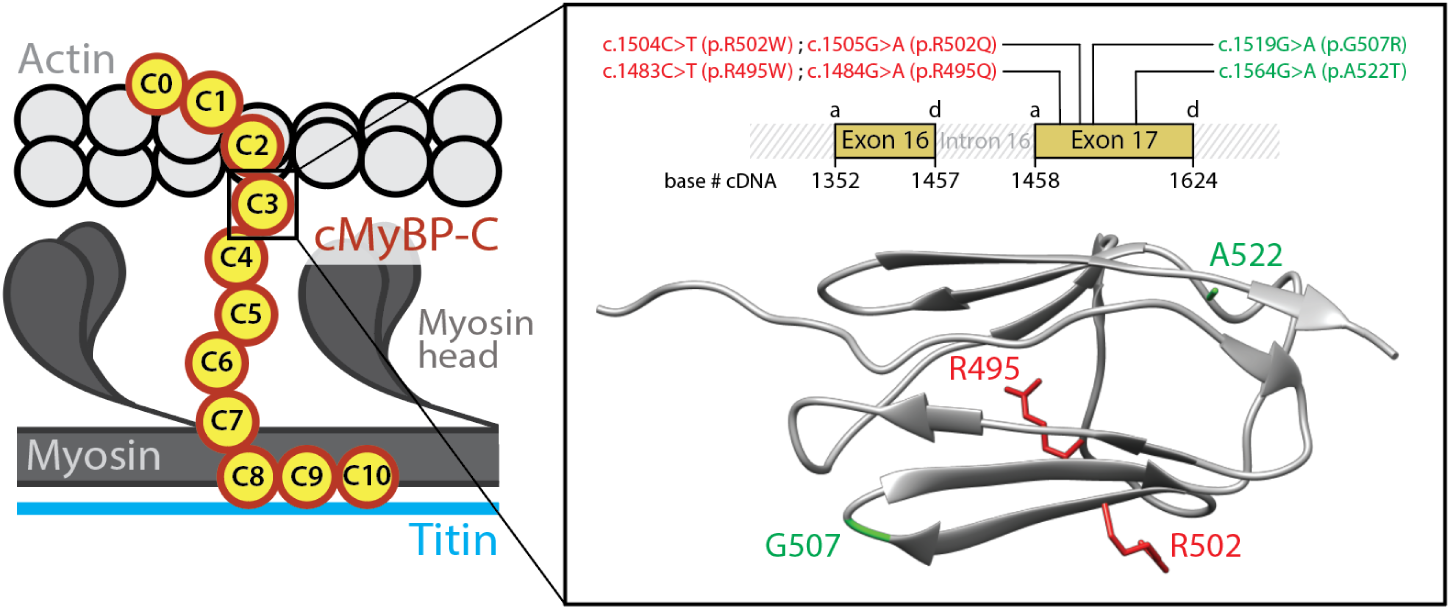
cMyBP-C variants tested in this report. The variants target the C3 central domain of cMyBP-C. *Inset*: the variants, which induce single nucleotide substitutions in *MYBPC3* exon 17, are presented using both cDNA and protein nomenclatures. Variants are colored according to their pathogenicity (red: pathogenic mutations; green: non-pathogenic variants). *MYBPC3* exons 16 and 17 codify for C3 domain and the position of their acceptor (a) and donor (d) splicing sites in the cDNA sequence is indicated. The ribbon diagram presents the Ig-like fold of the C3 domain, in which several β-strands arrange in a Greek key β-sandwich (pdb code 2mq0) ^27^. The side chains of the residues targeted by the variants are highlighted.

Preservation of normal RNA processing in R495Q and R502W mutants has been observed before using human myocardial biopsies ^31, 46^. In the case of mutation R502Q, no alteration of RNA splicing has been detected using the leukocyte fraction of human blood samples ^40^ and mini-gene constructs ^47^, two more readily available biological sources that provide results in excellent agreement with those obtained using myocardial samples ^40^. Using the former method, we studied RNA splicing of mutant R495W, and also sought additional validation that R495Q and R502W do not induce alteration of RNA splicing. We amplified by RT-PCR the region between exons 15 and 21 of *MYBPC3* mRNA and observed that amplification of wild-type (WT) and mutant samples results in bands at the ∼700 bp expected electrophoretic mobility (**Supplementary Figure S1a; Supplementary Text S1**). Preservation of canonical RNA splicing in the mutants was further confirmed by Sanger sequencing, which allows the detection of the variants in heterozygosis and the visualization of the correct 16/17 and 17/18 exon-exon boundaries (**Supplementary Figure S1b**).

To examine mutant protein stability, we did far-UV circular dichroism (CD) experiments using recombinant domains (**Supplementary File S1**; **Supplementary Figure S2a**). In agreement with previous reports, we found that WT and mutant domains have highly similar CD spectra showing a minimum at 215 nm, typical of β-structure-containing proteins ^27, 40, 48^ (**Supplementary Figure S3a**). The only remarkable difference was found in the spectrum of R502W, which displays lower CD signal at 230 nm (**Supplementary Figure S3b**). Since the high-resolution structures of C3 WT and R502W are very similar ^27^, we interpret this change as originating from the absorption of the extra tryptophan in the mutant ^49^. To explore if mutations induce structural destabilization, we examined the stability of the mutant domains at increasing temperatures by tracking the CD signal at 215 nm (**Supplementary Figure S4a**). The temperature at the midpoint of the denaturing transition, or melting temperature (T_m_), informs about the thermal stability of the domain. All mutations retain close-to-WT thermal stability (**Supplementary File S1**; **Supplementary Figure S4b**). The maximum drop in T_m_ was 5.3°C for mutant R502Q; however, this limited decrease in thermal stability can also be found in non-pathogenic missense variants targeting C3 and therefore cannot explain the pathogenicity of the mutation ^40^. In summary, none of the pathogenic variants studied in this report induce extensive protein destabilization.

### Mechanical destabilization in mutant cMyBP-C domains

The results in the previous section show that the selected pathogenic *MYBPC3* missense mutations preserve RNA splicing and protein thermal stability and thus it is unlikely that they cause HCM through classical protein haploinsufficiency. Most probably, these mutants are incorporated into the sarcomere but fail to provide proper functionality ^31, 50^. Since cMyBP-C tethers are subject to mechanical force in the sarcomere (**Figure 1b**), we used AFM to examine whether mutations alter the mechanical stability of the C3 domain (**Figure 3**). We first produced polyproteins consisting of eight repetitions of the WT or the mutant C3 domains (**Supplementary File S1**; **Supplementary Figure S2b**). Using AFM, these polyproteins were subject to 40 pN/s increasing pulling force while monitoring their length. Mechanical unfolding of a C3 domain within the polyprotein results in the extension of the polypeptide by 24-25 nm. The presence of multiple such unfolding steps fingerprints successful single-polyprotein recordings (**Figure 3a**) ^45^. We measured the force at which mechanical unfolding occurs for hundreds of WT and mutant domains, and built distributions of unfolding forces (**Figure 3b; Figure 4a; Supplementary Figure S5**). We found that the mean unfolding force (< ***F***_***u***_ >) of WT C3 domain is 90.6±0.3 pN, in agreement with previous measurements on cMyBP-C multidomain constructs ^24, 26^. All pathogenic and non-pathogenic variants induce slight mechanical destabilization under our experimental conditions, as indicated by lower mean unfolding forces (**Figure 3c**; **Table 1**).

**Table 1.** Nanomechanical properties of WT and mutant C3 domains. Errors are 83% confidence intervals of the fittings used to calculate the mechanical parameters. If these intervals do not overlap, differences are considered statistically significant (see Methods).

| Variant | Pathogenic variant | $\langle F_u \rangle$ (pN) | $r_0$ (s<sup>-1</sup>) | $\Delta x$ (nm) | $r_f$ (s<sup>-1</sup>) |
| --- | --- | --- | --- | --- | --- |
| R495Q | Yes | 89.0±0.6 | 0.009±0.001 | 0.25±0.01 | 0.28±0.14 |
| R495W | Yes | 87.7±2.4 | 0.020±0.004 | 0.21±0.01 | 0.11±0.03 |
| R502Q | Yes | 85.0±0.3 | 0.011±0.002 | 0.25±0.01 | 2.1±0.8 |
| R502W | Yes | 87.6±0.2 | 0.012±0.002 | 0.23±0.01 | 0.08±0.02 |
| G507R | No | 87.6±0.5 | 0.009±0.001 | 0.25±0.01 | 0.17±0.10 |
| A522T | No | 79.3±0.3 | 0.009±0.001 | 0.28±0.01 | 0.14±0.04 |
| WT | - | 90.6±0.3 | 0.012±0.002 | 0.22±0.01 | 0.16±0.06 |

**Figure 3.**
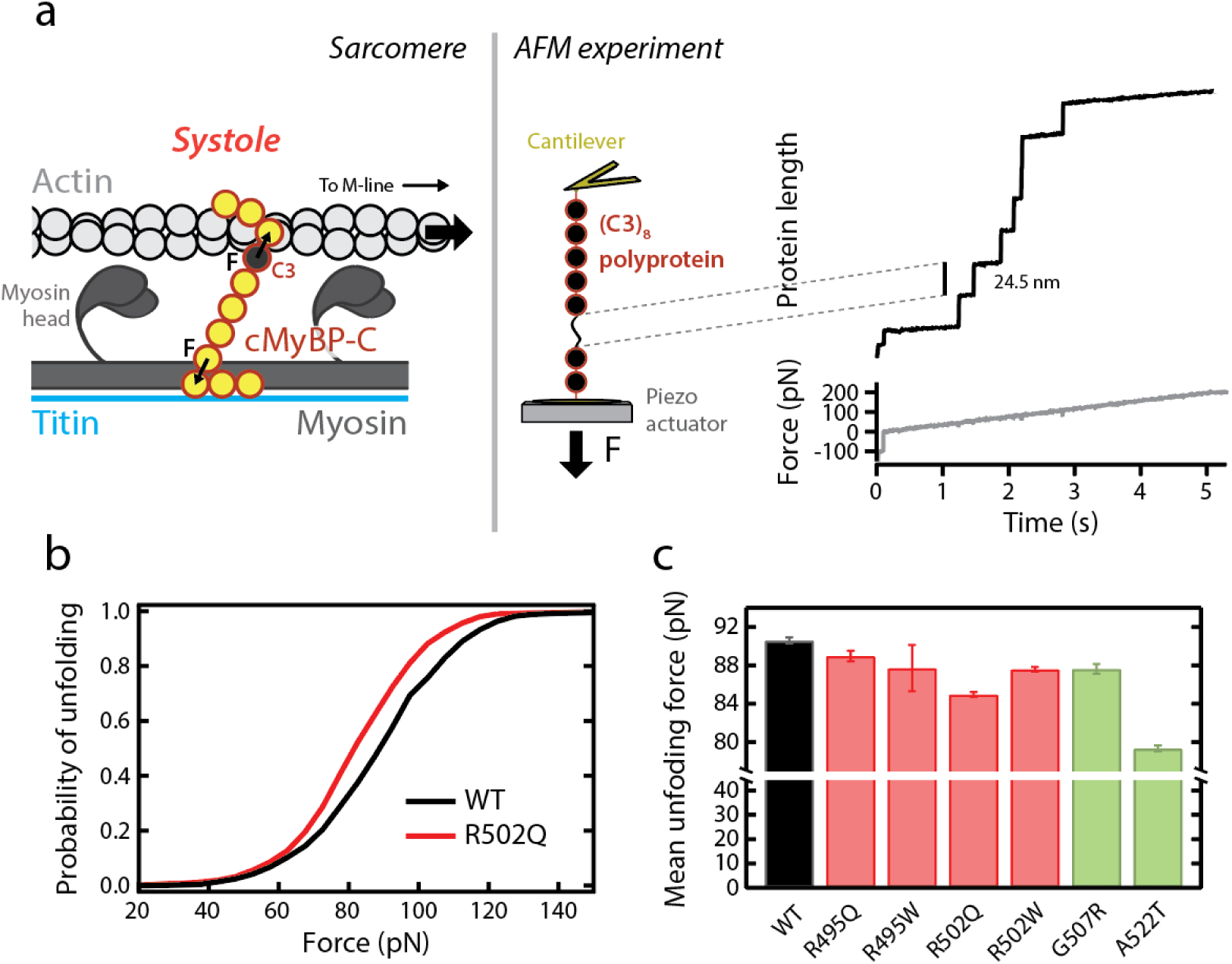
Characterization of the mechanical stability of WT and mutant C3 domains by single-molecule force-spectroscopy by AFM. **(a)** *Left*: cMyBP-C tethers experience end-to-end mechanical force during the contraction of actomyosin filaments in systole. The position of the C3 domain within cMyBP-C is indicated. *Right*: the mechanical properties of a (C3)8 polyprotein are measured using single-molecule AFM. In these experiments, a single polyprotein is tethered between a cantilever and a moving piezo actuator, and its length is recorded while a linear increase in force is applied. Unfolding events are detected as step increases in length of 24-25 nm ^45^. **(b)** Cumulative probability of unfolding with force for WT (n=1033 unfolding events) and R502Q domains (n=1254 unfolding events). **(c)** Mean unfolding forces, as obtained from Gaussian fits to distributions of unfolding forces (see also **Table 1**). Error bars correspond to 83% confidence intervals. Bars are colored according to the pathogenic status of the mutation (pathogenic, red; non-pathogenic, green).

**Figure 4.**
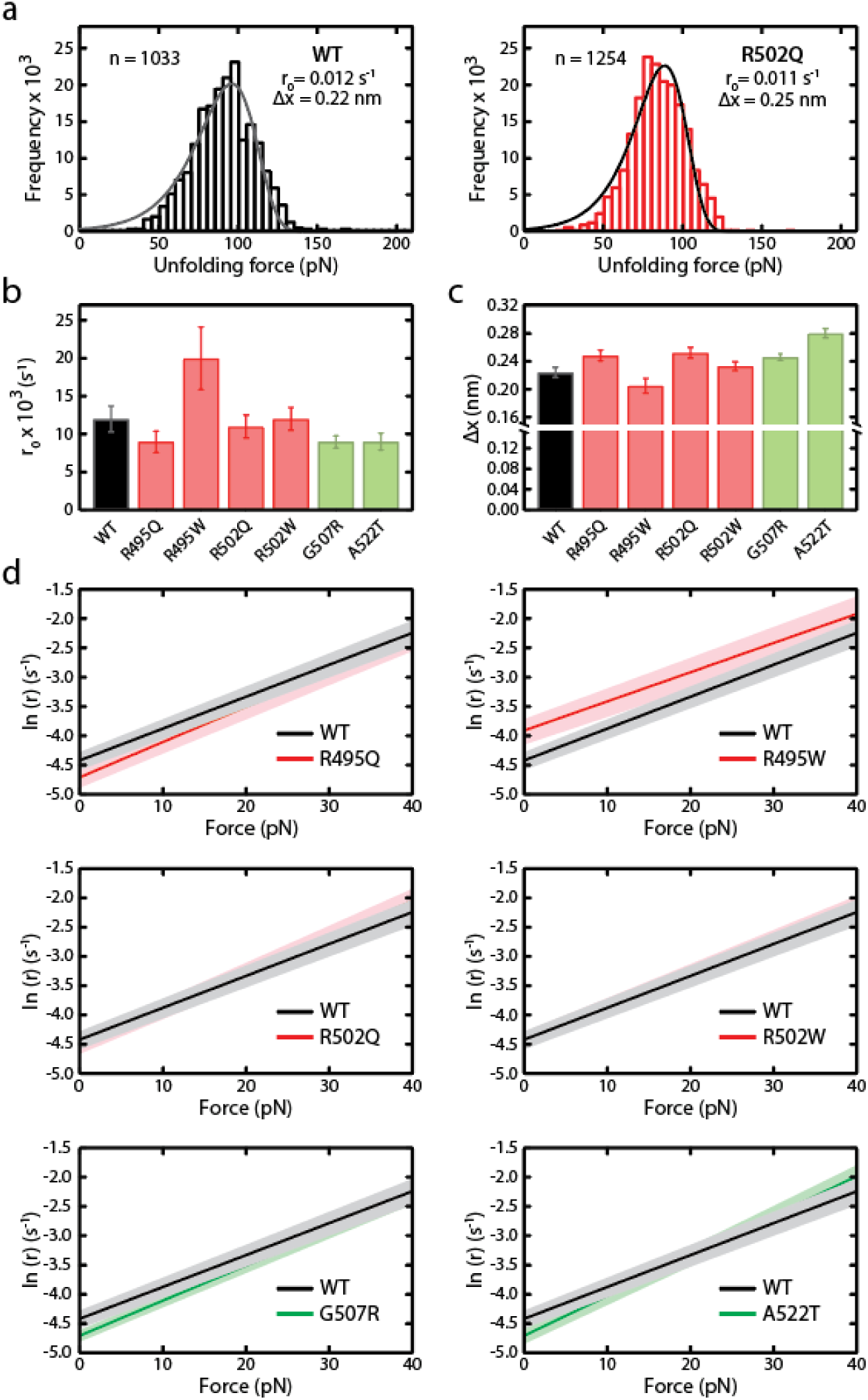
Characterization of *r*_0_ and Δ*x* parameters of WT and mutant C3 domains according to Bell’s model. **(a)** Distribution of unfolding forces obtained for WT and R502Q domains. Distributions were fit to Bell’s model ^51^ (fitting lines in black) and the resulting rate of unfolding at zero force, *r*_0_, and distance to the transition state, Δ*x*, are indicated. (**b,c**) *r*_0_ and Δ*x* values for WT and mutant C3 domains (see also **Supplementary Figure S5** and **Table 1**). Error bars correspond to 83% confidence intervals. Bars are colored according to the pathogenic status of the mutation (pathogenic, red; non-pathogenic, green). **(d)** Force dependency of rates of unfolding according to Bell’s model (red: pathogenic, green: non-pathogenic). 83% confidence intervals are indicated as shaded areas.

Experimental < ***F***_***u***_ > values are a consequence of the underlying free energy landscapes and the specific pulling conditions. Extrapolation of AFM data to alternative ranges of forces can be achieved using models that consider how the energy landscape is shaped by the applied mechanical force. We have done so by fitting our data to the Bell’s model (**Figure 4a; Supplementary Figure S5**) ^51-52^. According to this model, the rate of mechanical unfolding (*r*) is dependent on force (*F*) according to,

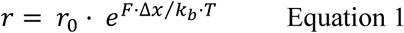

where *r*_0_ is the rate of unfolding at zero force, Δ*x* is the distance to the transition state of the mechanical unfolding reaction, *k*_*b*_ is the Boltzmann constant and *T* is the absolute temperature. Fits show that both pathogenic and non-pathogenic mutations can affect *r*_0_ and/or Δ*x*, therefore altering the mechanical behavior of C3 domains in a force-dependent manner (**Figure 4b,c**; **Table 1**). Using the parameters obtained from the fits, we estimated mechanical unfolding rates at low forces, which are challenging to probe experimentally using AFM but can be relevant in the context of cMyBP-C function in the sarcomere. This analysis showed that R495W domains unfold significantly faster than WT counterparts at forces below 40 pN, whereas the rest of the mutants behave very similarly to WT, including control, non-pathogenic variants (**Figure 4d**).

### Mechanical folding in missense mutants of cMyBP-C

To determine the ability of the different C3 domains to refold following mechanical unfolding, we did *unfolding-quench-probe* experiments (**Figure 5a**) ^54^. In these experiments, proteins are first pulled in a force ramp to high forces (*unfolding* pulse). Then, force is relaxed to 0 pN, at which domains can regain the folded state (*quench* pulse). Finally, in the *probe* pulse, the protein is pulled back to high forces. Unfolding steps in the *probe* pulse report on domains that refolded during the *quench* pulse. To obtain folding fractions, the number of unfolding events in the *probe* and *unfolding* pulses are compared. Folding rates were estimated by measuring folding fractions at different quench times. Compared to WT, mutant R502Q shows a 13x increased folding rate (*r*_*f*_). The remaining pathogenic mutations and non-pathogenic variants do not cause significant changes in the folding rate (**Figure 5b,c**; **Supplementary Figure S6; Table 1**).

**Figure 5.**
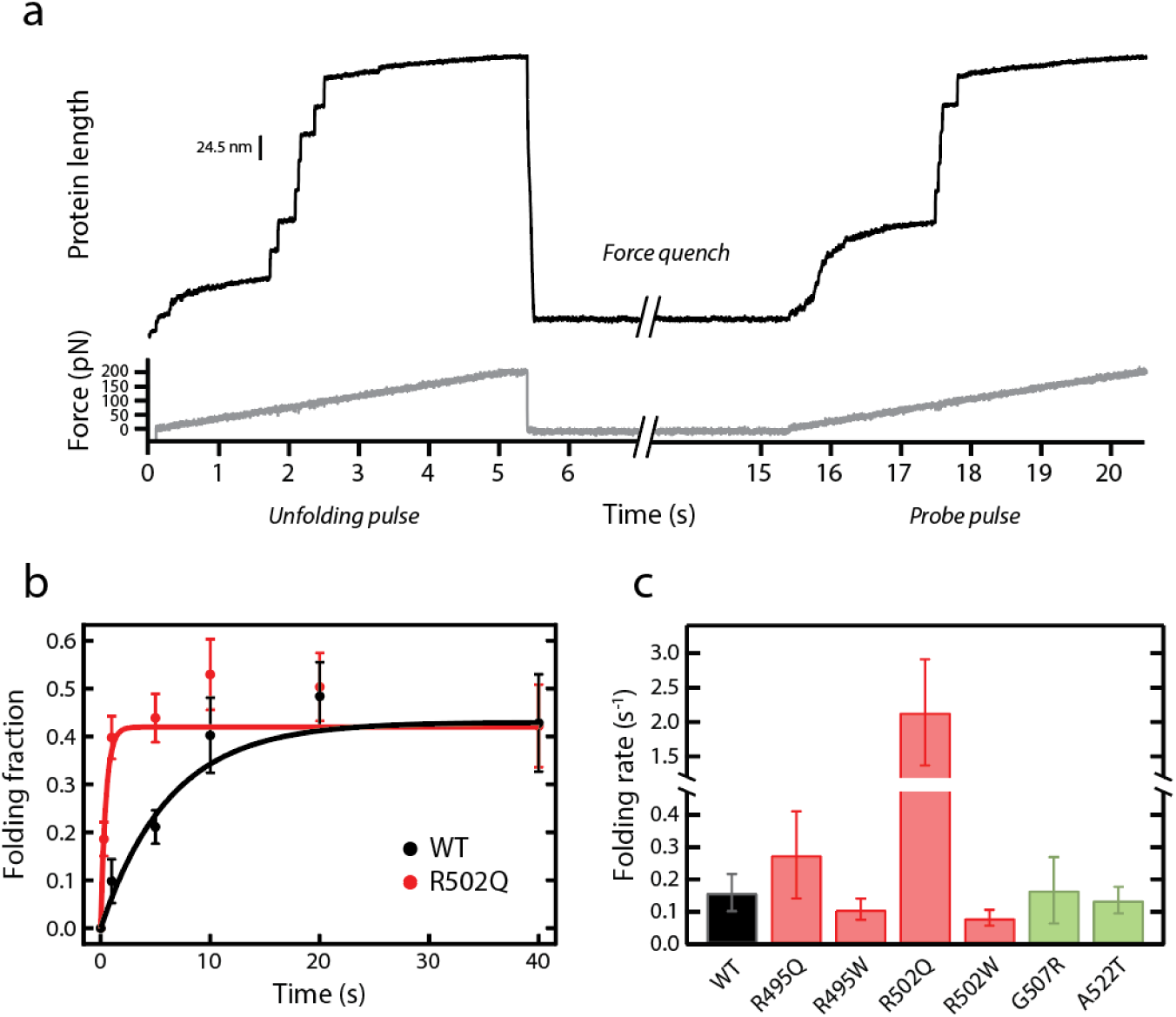
Characterization of mechanical folding of WT and mutant C3 domains. **(a)** Representative trace of mechanical refolding experiments by AFM. A single (C3)8 polyprotein is subject to an *unfolding* pulse, then force is quenched to 0 pN and finally the protein is pulled again to high forces in a *probe* pulse. Folding fractions are calculated comparing the number of unfolding events in the *probe* and the *unfolding* pulses. In the example shown, 5 out of 7 domains refolded during the *quench* pulse. **(b)** Folding fractions of C3 WT and R502Q at different quench times. Lines are exponential fits to the data. Error bars are standard errors of the mean estimated by bootstrapping ^53^ (n≥56 and n≥86 unfolding events for all WT and R502Q data points, respectively). **(c)** Mechanical folding rates for C3 WT and its mutants, obtained from exponential fits to refolding data (see also **Supplementary Figure S6** and **Table 1**). Error bars are 83% confidence intervals. Bars are colored according to the pathogenic status of the mutation (pathogenic, red; non-pathogenic, green).

## DISCUSSION

It has been proposed that HCM mutations lead to increased sarcomeric output and myocyte hypercontractility ^55-58^. Indeed, mavacamten, an inhibitor of myosin motor activity, is able to prevent development of HCM in mouse models ^59^ and also shows beneficial effects in clinical trials ^7^. Many HCM-causing mutations in myosin affect directly the mechanochemical cycle of the protein resulting in enhanced force generation ^60-61^. Interestingly, the proportion of low-activity SRX myosin heads is decreased in HCM myocytes devoid of cMyBP-C but is normalized upon treatment with mavacamten ^57, 62-63^. The emerging unifying view is that, regardless of the specific mutation, HCM is primarily a mechanical disease at the molecular level; however, it remains unknown how missense mutations in the central region of cMyBP-C can lead to hypercontractility ^64^. In this work we have studied several pathogenic mutations targeting the central C3 domain of cMyBP-C, the region of the protein with the highest number of HCM-causing missense mutations ^13^. We verified that our variants do not induce classical protein haploinsufficiency drivers specifically associated with other HCM mutations (**Supplementary Figure S1, S3, S4**) ^40^. Hence, these variants trigger HCM by yet-to-identify molecular mechanisms.

The modulatory mechanisms of cMyBP-C on contraction are complex and far from being completely understood. While the C8-C10 C-terminal domains of cMyBP-C play a structural role providing strong anchorage to the thick filament, the C0-C2 N-terminal fragments can bind both actin filaments and myosin globular heads resulting in sophisticated control of their activity through direct mechanical load (**Figure 1b**) or via conformational changes that are dependent on phosphorylation and calcium levels ^15^. In this highly intricate regulatory landscape, several possibilities can be envisioned by which altered cMyBP-C nanomechanics, as detected here for R495W and R502Q (**Figures 4d, 5c**), can perturb sarcomere function.

Let us first consider a purely mechanical scenario. Current models on cMyBP-C modulation of sarcomere contraction state that the central region of the protein is subject to end-to-end mechanical force that results in a viscous drag opposing contraction (**Figure 1b**) ^15, 21, 23-24, 65-66^. This drag force depends on cMyBP-C stiffness. If we assume that 5 cMyBP-C domains bridge radially the thick and thin filaments ^16^, using the freely jointed chain (FJC) model of polymer elasticity ^67^, we can predict that the force experienced by cMyBP-C tethers increases by >250% during a 10-nm myosin power stroke (4.4 nm contour length, Lc, per domain, and 20 nm Kuhn length, Lk) ^68^ (**Figure 6**). This force is expected to decrease to less than pre-power-stroke values if a cMyBP-C domain mechanically unfolds in the process (Lc = 0.4 nm per amino acid, 90 amino acid domain size, Lk = 1.32 nm for unfolded polypeptide regions) ^68^. According to our AFM data, this low-force state is up to 66% more frequent in mutant R495W (**Figure 4d**), which is expected to reduce the average viscous load generated by mutated cMyBP-C. We speculate that the higher speed of folding detected for R502Q could alter other steps of the mechanochemical cycle of actomyosin filaments, for instance by generating more mechanical work during relaxation ^69^. It is important to stress that force estimates in **Figure 6** are highly dependent on the specific geometry of cMyBP-C tethers and on the mechanical parameters considered. Although the model exemplifies how cMyBP-C’s viscous load can be reduced by mechanically labile domains, the exact forces experienced by cMyBP-C in the sarcomere, which so far remain unknown, may differ from the values shown in **Figure 6**.

**Figure 6.**
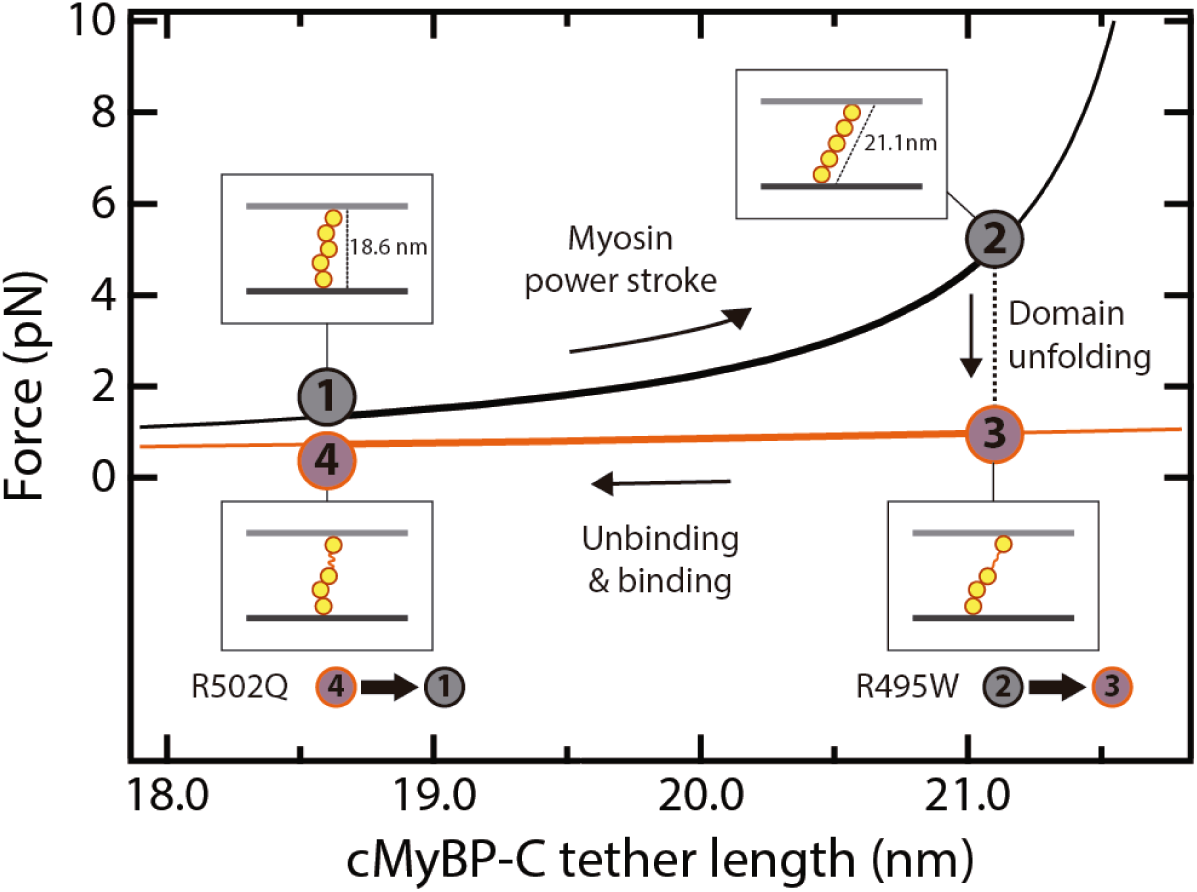
Model of mechanical modulation by cMyBP-C tethers and the influence of domain unfolding. FJC-estimated increase in force generated by a fully folded cMyBP-C tether (black) during a myosin power stroke. If one of the domains of cMyBP-C unfolds, force is reduced (orange). The model considers a radial distribution of domains C3-C7 ^16^, and that anchoring C2 and C8 domains (not shown for simplicity) are located at the center of the thin and thick filaments, respectively, so that they contribute half their diameter to bridge the 23-nm interfilament space ^20^. The increase in cMyBP-C length during a myosin power stroke was estimated using the Pythagorean theorem considering a 10 nm power stroke ^12^. In the graph, we also considered that cMyBP-C can dissociate from actin sites at a slower rate than that of myosin power strokes. Mutation R495W increases the rate of mechanical unfolding of C3, whereas R502W leads to faster C3 folding at low forces.

The interaction of cMyBP-C terminal domains with several sarcomeric proteins, resulting in modulation of their activity, is well documented ^15^. The possibility exists that the central domains of cMyBP-C transitorily participate in those binding reactions, or have yet unidentified binding partners. Hence, as described in mutants targeting terminal cMyBP-C domains ^36, 70-71^, mutations in central domains of cMyBP-C could also interfere with binding reactions either directly or allosterically. Interestingly, the interactome of proteins under force, such as titin and talin, is dependent on mechanical load ^54, 72^. Hence, the mechanical unfolding of cMyBP-C central domains could result in the exposure of cryptic binding sites. In this scenario, altered mutant cMyBP-C nanomechanics would also cause perturbations of the interaction landscape of cMyBP-C.

Truncating mutations in cMyBP-C that cause HCM result in normal levels of mutated mRNA but no detectable truncated polypeptide in the myocardium, probably due to its degradation by cellular protein quality control systems, such as the ubiquitin/proteasome system (UPS). Beyond insufficient mechanical modulation in cMyBP-C deficient myocardium, the exhaustion of these protein control systems can also contribute to the pathogenesis of HCM ^73-76^. Since mechanical destabilization results in increased protein unfolding, a well-known trigger of the UPS ^77^, it is possible that nanomechanical destabilization of cMyBP-C missense mutants can result in activation of protein quality control systems.

In summary, our single-molecule experiments reveal that pathogenic missense mutations in cMyBP-C can induce mechanical destabilization and alterations in the mechanical folding properties of the targeted domain. We propose that these nanomechanical phenotypes, which are not found in non-pathogenic variants, can perturb the function of cMyBP-C in the sarcomere by several mechanisms, potentially contributing to HCM pathogenesis. Similar nanomechanical phenotypes may be also found in pathogenic mutations targeting other proteins with mechanical roles, such as titin ^78^, talin ^79^, filamin ^80^, lamin ^81^, and α-catenin ^82^. Future studies will investigate the prevalence of nanomechanical phenotypes in other cMyBP-C mutations, and their impact in low-force transitions that are now amenable for experimental observation ^83^.

## METHODS

### Human samples

The procurement of human samples was achieved following the principles outlined in the Declaration of Helsinki according to a project approved by the *Comité de Ética de Investigación* of *Instituto de Salud Carlos III* (PI 39_2017) and by the Ethics Committee of the Naples University Federico II “Carlo Romano” (Protocol number 157/13).

### Analysis of RNA splicing

Leukocytary fractions from carrier’s venous blood were treated with Trizol (Thermo Fisher Scientific) to extract total RNA. Then, cDNA was obtained by retro-transcription of total RNA with random primers and Superscript™ IV VILO™ Master Mix (Thermo Fisher Scientific). To amplify the region spanning *MYBPC3* exons 15 through 21, we used primers MyBPC Forward (5’-CAAGCGTACCCTGACCATCA-3’) and MyBPC Reverse (5’-GGATCTTGGGAGGTTCCTGC-3’). To avoid amplification of genomic DNA, the annealing region of the reverse primer targets the exon 20/21 junction **(Supplementary Text S1**). The resulting PCR amplification products were purified using the Qiaquick PCR purification Kit (Qiagen). Finally, the purified fragments were sequenced using the Sanger method. We consider that standard RNA processing is not altered when electropherograms allow the unambiguous reading of the expected WT *MYBPC3* cDNA sequence at the junction between exons 16, 17 and 18. Since patients carry mutations in heterozygosis, a double peak corresponding to the WT and mutated base was detected at the variant position, as expected.

### Protein expression and purification

The production and purification of the (C3)_8_ polyproteins was done as reported before ^45^, except for the case of non-pathogenic variants, which were also expressed by overnight induction of BLR(DE3) *E*.*coli* cultures with 0.4 mM isopropyl β-d-1-thiogalactopyranoside, at 18°C and 250 rpm agitation. Mutations were introduced by PCR. Polyprotein-coding mutated cDNAs were obtained by an iterative cloning strategy using BamHI, BglII and KpnI restriction enzymes, as described ^84-85^. To produce monomeric C3 domains, the corresponding cDNA was cloned in a custom-modified pQE80L plasmid using BamHI and BglII restriction enzymes. Sanger sequencing was performed on all final expression plasmids. Monomers and mutant polyproteins were purified following the same protocol as for wild-type (C3)_8_ ^45^, which includes two rounds of purification using nickel-based affinity and size-exclusion chromatographies. In this final step, elution was carried out in 10 mM Hepes, pH 7.2, 150 mM NaCl,1 mM EDTA for polyproteins, while monomers were recovered in 20 mM NaPi, pH 6.5 and 63.6 mM NaCl. Proteins were stored at 4°C. SDS-PAGE analysis was carried out to evaluate the isolating process and to identify the purest and highest concentrated fractions.

### Circular dichroism

CD experiments were conducted in a Jasco J-810 spectropolarimeter. Protein fractions at 0.1-0.5 mg/mL were tested in 20 mM NaPi, pH 6.5 and 63.6 mM NaCl. CD signal was recorded every 0.2 nm at a speed of 50 nm/min. Data were collected with standard sensitivity (100 mdeg). Four different scans were performed for each construct, which were later averaged to obtain the final spectra. The absorbance contribution of the buffer (baseline spectrum) was subtracted from the protein spectra, which were finally normalized according to the protein concentration. Protein concentrations were estimated from A_280_ measurements considering theoretical extinction coefficients according to ProtParam Tool (**Supplementary File S1**) ^86^. To examine thermal stability, protein samples were heated at 30°C/h from 25 to 85°C using a Peltier temperature controller while recording CD signal at 215 nm (0.5°C data pitch). The recorded CD signal changes as the protein denatures and unfolds, and the melting temperature (T_m_) was calculated by performing a sigmoidal fitting to the denaturing curves using IGOR Pro (Wavemetrics).

### Single-molecule atomic force spectroscopy

Single-molecule AFM experiments were done in an AFS force-clamp spectrometer (Luigs & Neumann) ^41^. 1-20 uL of a 0.04-1.5 mg/mL solution of the purified polyprotein in 10 mM Hepes, pH 7.2, 150 mM NaCl, 1 mM EDTA buffer were deposited onto a gold-coated cover slip (Luigs & Neumann). We used silicon nitride MLCT cantilevers (Bruker AFM Probes), with reflective 60-nm gold coating on their back side. These cantilevers were calibrated using the thermal fluctuations method ^87^. Typical spring constants were in the range of 15-20 pN/nm. Single polyproteins were picked up by pressing the cantilever onto the gold surface at a contact force of 500-2000 pN for 0.8-2 s, and subject to 40 pN/s linear increase in force until their detachment. During this stretching, the length of the polyprotein is measured and unfolding events are detected as specific stepwise increases in extension. The mechanical unfolding of C3 domains results in steps of 24-25 nm; only traces containing two or more unfolding events and with detachment forces >175 pN were analyzed ^45^. Traces where the fingerprint was interrupted by unidentifiable events were discarded. Unfolding forces were recorded in at least three different experiments performed with different cantilevers. Only experiments with low laser interference (peak-to-peak height in baseline force-extension traces lower than 25 pN) were included in the analysis of unfolding forces ^41^. Mean unfolding forces were obtained from Gaussian fits to histograms of unfolding forces. Some of the unfolding data for wild-type C3 have also been included in a technical report ^45^. Distributions were also fit to the Bell’s model of protein unfolding under a force ramp to get the value of the unfolding rate at zero force, *r*_0_, and the distance to the transition state, Δ*x* ^51^. For folding experiments, we programmed a force regime consisting of two ramps separated by a *quench* pulse. To quantify folding rates, we calculated folding fractions at different quench times. The bootstrap method was used to estimate standard errors of the mean for the folding fractions ^53^. Time courses of folding were fit to an exponential function,

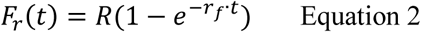

where *F*_*r*_*(t)* is the folding fraction as a function of time, *R* if the maximum folding fraction and *r*_*f*_ is the folding rate. Fits considered that the folding reaction was complete at 40 s. Values of *R*<1 in the folding of Ig domains are common and are due to misfolded states that are highly dependent on experimental conditions ^54, 88^. Hence, we focused on folding rates, which is a more robust molecular parameter. Analysis was done in IGOR Pro (Wavemetrics).

### Statistics

Statistical significance of the differences in parameters between WT and mutant domains were inferred from 83% confidence intervals, which were estimated using IGOR Pro (**Supplementary File S1, Table 1**). No overlapping intervals suggest that the null hypothesis can be rejected with p<0.05 ^89-90^.

**Supplementary Information** is available in the online version of the article. Supplementary info includes 1 Supplementary File, 1 Supplementary Text, 6 Supplementary Figures, and Supplementary References.

## Acknowledgements

JAC acknowledges funding from the *Ministerio de Ciencia e Innovación* (MCIN) through grants BIO2014-54768-P, BIO2017-83640-P (AEI/FEDER, UE), EIN2019-102966, RYC-2014-16604, and BFU2017-90692-REDT, the European Research Area Network on Cardiovascular Diseases (ERA-CVD/ISCIII, MINOTAUR, AC16/00045) and the *Comunidad de Madrid* (consortium Tec4Bio-CM, S2018/NMT-4443, FEDER). The CNIC is supported by the *Instituto de Salud Carlos III* (ISCIII), MCIN and the Pro CNIC Foundation, and is a Severo Ochoa Center of Excellence (SEV-2015-0505). We acknowledge funding from ISCIII to the *Centro de Investigación Biomédica en Red* (CIBERCV), CB16/11/00425. CSC is the recipient of an FPI-SO predoctoral fellowship BES-2016-076638. MRP was the recipient of a PhD fellowship from the Italian Ministry of Education, Universities and Research (MIUR). CPL was a recipient of CNIC Master Fellowship. We thank Natalia Vicente for excellent technical support (through grant PEJ16/MED/TL-1593 from *Consejería de Educación, Juventud y Deporte de la Comunidad de Madrid* and the European Social Fund). We thank the Spectroscopy and Nuclear Magnetic Resonance Core Unit at CNIO for access to CD instrumentation. We thank Andrea Thompson and Sharlene Day for their insights. We thank all members of the *Molecular Mechanics of the Cardiovascular System* team for helpful discussions.

## Author contribution

JAC conceived the project. DSO, SV, FD, GF, PGP and JAC ensured procurement of human samples. MRP did experimental analysis of RNA splicing. CSC and DVC cloned and purified proteins. CSC and EHG did circular dichroism experiments. CSC, CPL, IUI and JAC did single-molecule AFM experiments. CSC, MRP, DSO, LM, EHG and JAC interpreted data in the context of the available literature. CSC and JAC drafted the manuscript with input from all authors.

