## Supplementary Information for "Nanomechanical phenotypes in cMyBP-C mutants that cause hypertrophic cardiomyopathy"

Supplementary Information includes:

- **Supplementary File S1 (separate excel file)**
- **Supplementary Text S1**
- **Supplementary Figures S1-S6**
- **Supplementary References**

**Supplementary Text S1. Genomic region of *MYBPC3* (ENST00000545968.6) showing the location of the missense variants studied in this report.** The positions targeted by the variants are highlighted in bold and colored according to the pathogenic status of the substitution (red: pathogenic, green: non-pathogenic). The hybridization sequences of the MyBPC Forward and Reverse oligos used in the experimental characterization of RNA splicing are double underlined. MyBPC Reverse primer hybridizes in the junction between exons 20 and 21 to avoid amplification of gDNA. Exons are shaded in grey.

| | | |
|---|---|---|
| **Exon 15** | cacccagGTACATCTTTGAGTCCATCGGTGCCAAGCGTACCCTGACCATC | MyBPC Forward |
| | AGCCAGTGCTCATTGGCGGACGACGCAGCCTACCAGTGCGTGGTGGGTGG | |
| | CGAGAAGTGTAGCACGGAGCTCTTTGTGAAAGgtgggcctgggacctgag | |
| | gatgtgggaacctggggaggagatggcctcaggggagccaaccctcatgc | |
| **Exon 16** | tcaccctgcctggacagAGCCCCCTGTGCTCATCACGCGCCCCTTGGAGG | |
| | ACCAGCTGGTGATGGTGGGGCAGCGGGTGGAGTTTGAGTGTGAAGTATCG | |
| | GAGGAGGGGGCGCAAGTCAAATGgtgagttccagaagcacggggcatggg | |
| | tgttgggggcatctgcccagaagaggccacagcacttgccacccacccac | |
| **Exon 17** | ccggctagGCTGAAGGACGGGGTGGAGCTGACC**CG**GGAGGAGACCTTCAA | c.1483C>T (p.R495W); c.1484G>A (p.R495Q) |
| | ATAC**CG**GTTCAAGAAGGAC**G**GGCAGAGACACCACCTGATCATCAACGAGG | c.1504C>T (p.R502W); c.1505G>A (p.R502Q) |
| | CCATGCTGGAGGAC**G**CGGGGCACTATGCACTGTGCACTAGCGGGGGCCAG | c.1519G>A (p.G507R) |
| | GCGCTGGCTGAGCTCATTGTGCAGGgtgagcctggctgggggggcacatg | c.1564G>A (p.A522T) |
| | aggctttagggcttggacccctcagcccccaccccacctagccctgtagg | |
| | ggagcaggcaaacctgggctcaaggcctctgtgaccttgggcctttgaca | |
| | gcctaaacctcatccttctcatctctaaaatgacgatgctgaggtgaccg | |
| | ctccagagggtactgagagggtcatacgggcctggtgagcatcaccaccc | |
| | ctcagacacttgaggttccttatgtgcactgcaccatcacaggcaaacat | |
| | ggcacacacaggcacacgtgttttcacatgcccacatgcacccagacacg | |
| | tgcgcaccagcgcccatgggcccacacgccctccacagggattcacgcca | |
| **Exon 18** | cacccacacaccctcccgagctcaatggctctgccctgccctgcagAAAA | |
| | GAAGCTGGAGGTGTACCAGAGCATCGCAGACCTGATGGTGGGCGCAAAGG | |
| | ACCAGGCGGTGTTCAAATGTGAGGTCTCAGATGAGAATGTTCGGGGTGTG | |
| | TGGCTGAAGAATGGGAAGGAGCTGGTGCCCGACAGCCGCATAAAGGTGTC | |
| | CCACATCGGGCGgtgagtgtgcagggcaggtggatgggacaggtggagac | |
| | tgagatggagacagagagagacacagggagaaacagagagagagagataa | |
| | agacaggtggacagagaggcagatgcacagagacaggacagagactgaga | |
| | cagagacccacagagagggagaaacagacagacccaaaaagacaaagaga | |
| | cagaaagacagaaatagttctatagagacagaaagatatgcacagggagg | |
| | cacacacataggagagttcgtcagagacagagaagaacagagggacagag | |
| | acaaagaggcagcagggagagacacggacggggtcgagcgctccccagca | |
| | ccctcccccagcttgtgaccagggcagggcaggaggccaccaaggaggcc | |
| | ggctctggccagaggccctcactgcctctcccgccctcttgctcatccct | |
| | ggagcaactgggctggccgtgctgggcccaagttcacagagattaatgag | |
| | acccagcccctgcccagtctccaagggcccagccaaggagatgctttgat | |
| | gtagaaatcacttttacgctgcctacctctgaaggcacattaaaaagctg | |
| | gaaaattagataatatagaaaaaaaaaggcatttttaaaaatcagaatac | |
| | caacaagccaggacaaggtgagggcctcctgggaccctggctggggtatc | |
| | tggcaaggccaggggtgtgtggcccagtggggtcccctgagccactgctc | |
| **Exon 19** | ccctgcagGGTCCACAAACTGACCATTGACGACGTCACACCTGCCGACGA | |
| | GGCTGACTACAGCTTTGTGCCCGAGGGCTTCGCCTGCAACCTGTCAGCCA | |
| | AGCTCCACTTCATGGgtgagcctgctccagggtagggtgggtcggggcag | |
| | tggggccaggagcccctgtcactgggccctgcctttgccccgtgctact | |
| **Exon 20** | tgctcttccttctcttgcagAGGTCAAGATTGACTTCGTACCCAGGCAGG | MyBPC Reverse |
| | gtaagtcttggggcccctgagtcttggttctgctcactttcccgcaaccc | |
| | acccacgcaggcctcccctccactcacagagggaagagcaaggatcaaaa | |
| | gtgggggtcctggggcccagcttcagctttgccactcatgaccatgggtc | |
| | aattataatctctgtgagcctcagtttcctcttctctaaaatgggaaccc | |
| | tgatagtgccacctcataggatagttataaagatgaggtgacatgatgta | |

tcagacctgtcaagtgctgtcacatgtgtgactgttattattacatcttt
tcttcttcttcctttttttttattttattttttttgagacggagtctcc
ctctgtcgcccacgctggagagaagtggcgcgatctcggctcactgcaag
ctccgcctcccaggttcacaccactctcctgcctcagcctcctgagtagc
tgggactacaggcgcccgccaccatgcccagctaattatttttttgtattt
ttagtagagacggggtttcaccatgttagccaggatggtttcgatctcct
gatctcatgatctgcctgcctcggcctcccaaagtgctgggattacaggc
gtgagccactgcgtccagactgctttgctttgtttaactgagggcagatt
cctgatttatttggccagggaaccttttttaatgtgtgcatgtgtgcgtgc
atgtgtgcgcgcgtgtgtgtgtgtgcatgtgtctgtgtgtgtatgtatat
atgtgtgtctgtgtctgtgtgtgtgcgtgtatgtatgtgtatttgtgtct
gtgtgcatgtgtgtgtctgtgtgtgcatgtgtgtatgtgtatgtgtgtct
gtgtgtatgtgtatatgcgtgtgtgtgtgtgtattgtgtgtgtgtgtatg
tgtgtggcaagacactgatgttcccccagttttcaaaatgctatgctagg
tgacctgacctgaatattacaagcctcccagccaacgtcctggggtgatg
ggaatgttccagatcttggttggattaataattgcatgggtgtattcatc
tgtcagaactcagcctactgtcagatctgagcataaactatacctcaata
caaatctaaaaaaccaataaccagggggcccaggctcttgagcagccccgc
cccagtgacctgtgctcctcctggctctcccgtttctctgaactacattg

**Exon 21** tgtcttctgcagAACCTCCCAAGATCCACCTGGACTGCCCAGGCCGCATA MyBPC Reverse (continued)
CCAGACACCATTGTGGTTGTAGCTGGAAATAAGCTACGTCTGGACGTCCC
TATCTCTGGGGACCCTGCTCCCACTGTGATCTGGCAGAAGGCTATCACGCAG

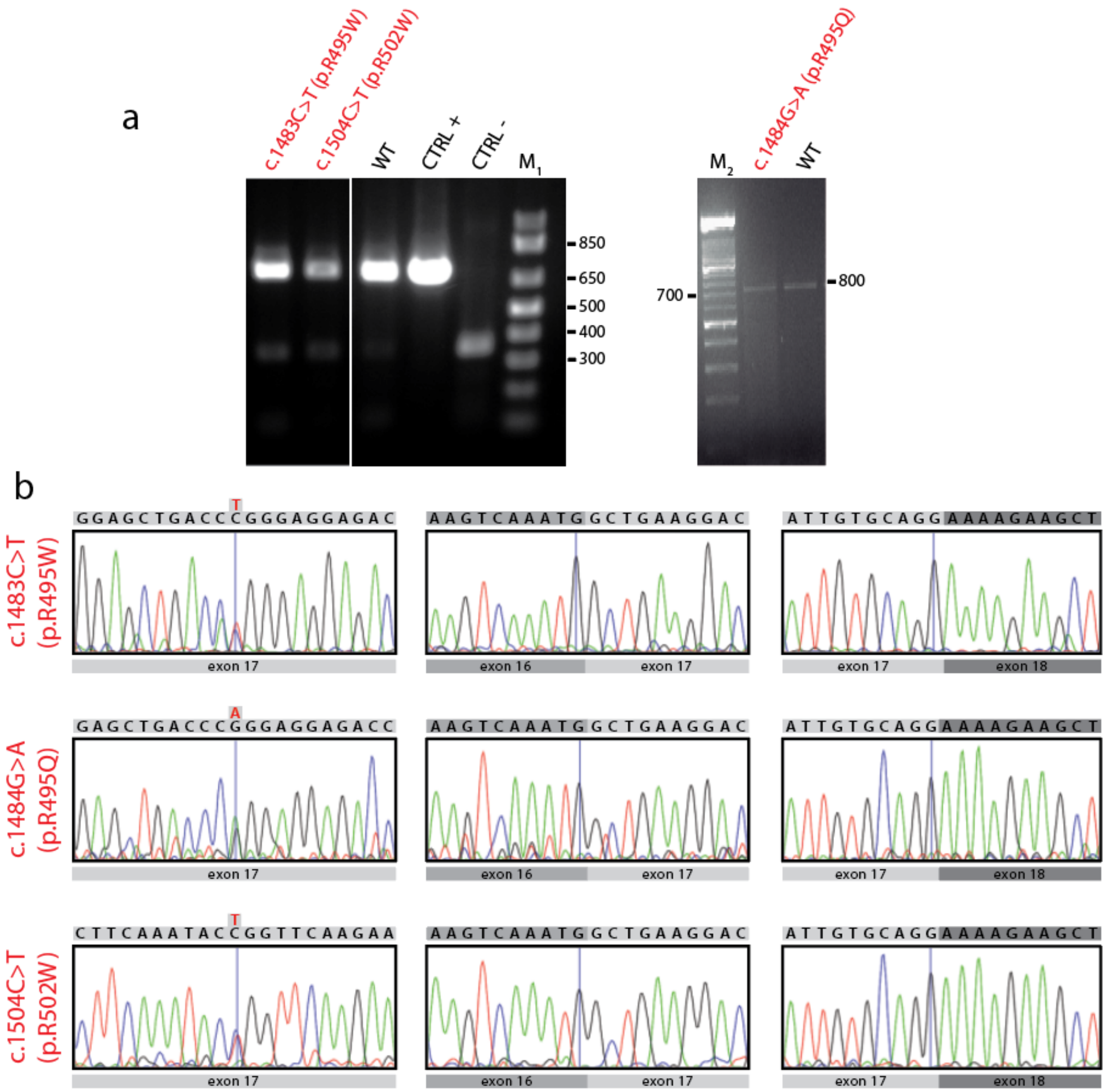


**Supplementary Figure S1. Examination of RNA splicing in the pathogenic *MYBPC3* missense mutations. (a)** RT-PCR analysis of mRNA isolated from peripheral blood of carriers. CTRL + corresponds to the analysis of mRNA obtained from healthy myocardium, whereas CTRL – refers to the transcript evaluation from HeLa cells, which do not express *MYBPC3*. The examination of mRNA obtained from the peripheral blood of healthy individuals is labelled as WT. Samples c.1483C>T (p.R495W), c.1504C>T (p.R502W), WT, CTRL+ and CTRL- were all loaded in the same agarose gel together with other variants reported elsewhere [1]. $M_1$: 1 Kb plus DNA ladder (Invitrogen). $M_2$: DNA Molecular Weight Marker XIV (Roche). Base pairs are indicated for both markers. **(b)** Sanger sequencing results, showing the presence of the variants in heterozygosis (*left*) and normal 16/17 (*middle*) and 17/18 (*right*) exon/exon boundaries. Canonical and mutated sequences are indicated on top of the Sanger electropherograms.

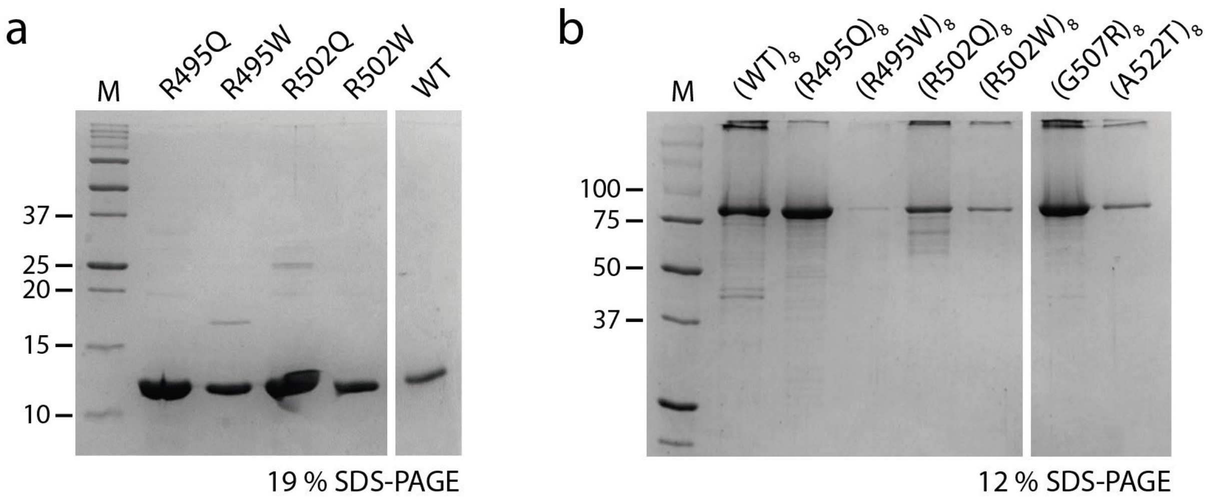


**Supplementary Figure S2. SDS-PAGE analysis of the purified proteins used in this report. (a)** Monomeric WT and mutant C3 domains used for circular dichroism experiments. **(b)** WT and mutant $(C3)_8$ polyproteins used for single-molecule AFM mechanical characterization. M: Precision Plus Protein Unstained Standard (Bio-Rad). Molecular weights in kDa are indicated.

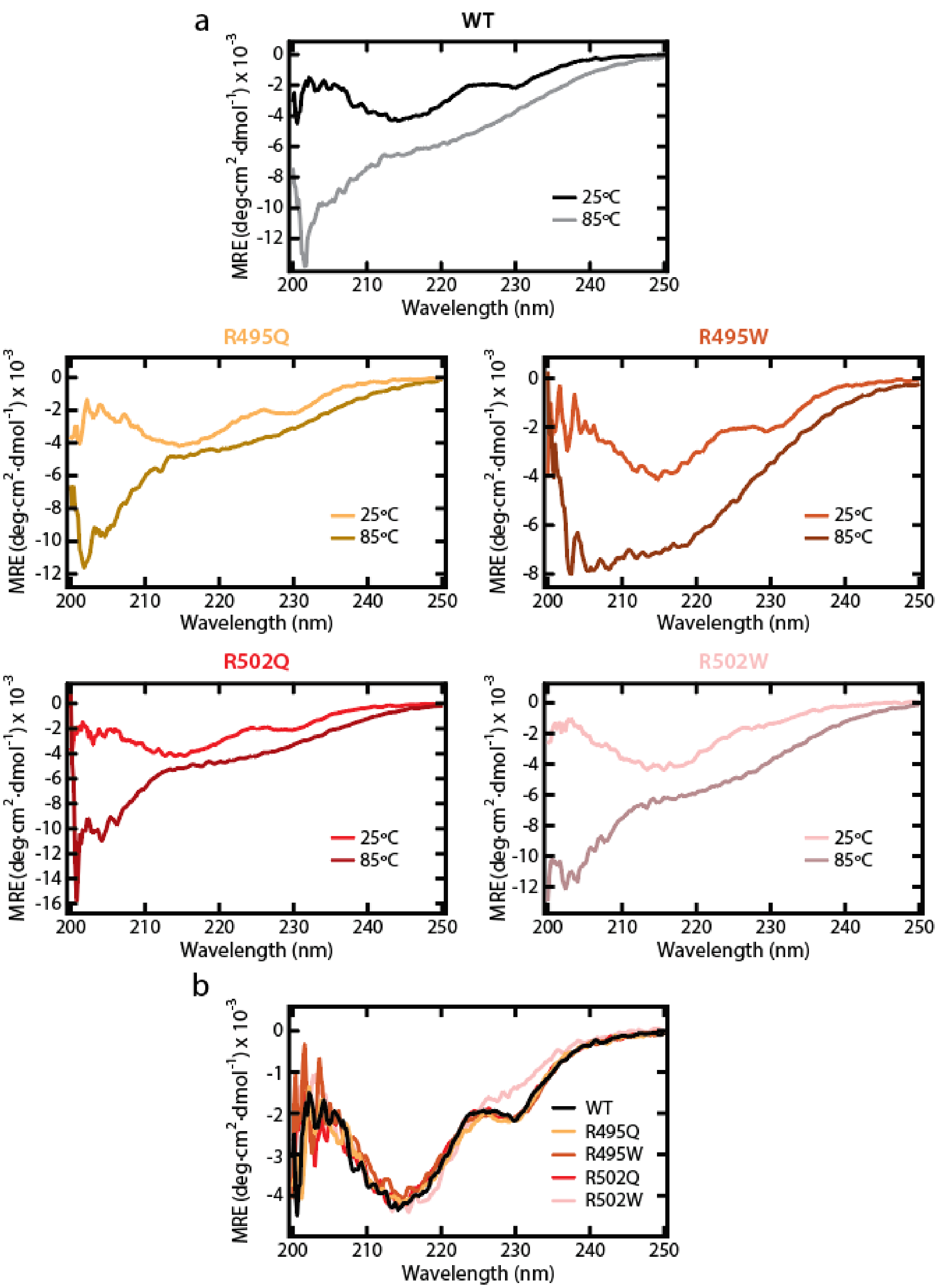


**Supplementary Figure S3. Structural characterization of WT and mutant C3 domains by circular dichroism (CD). (a)** Far-UV CD spectra of recombinant WT and mutant C3 monomers were recorded at 25ºC and 85ºC. These same spectra for WT has been presented before in [1]. **(b)** Comparison of CD spectra obtained at 25ºC for the WT and mutant C3 domains.

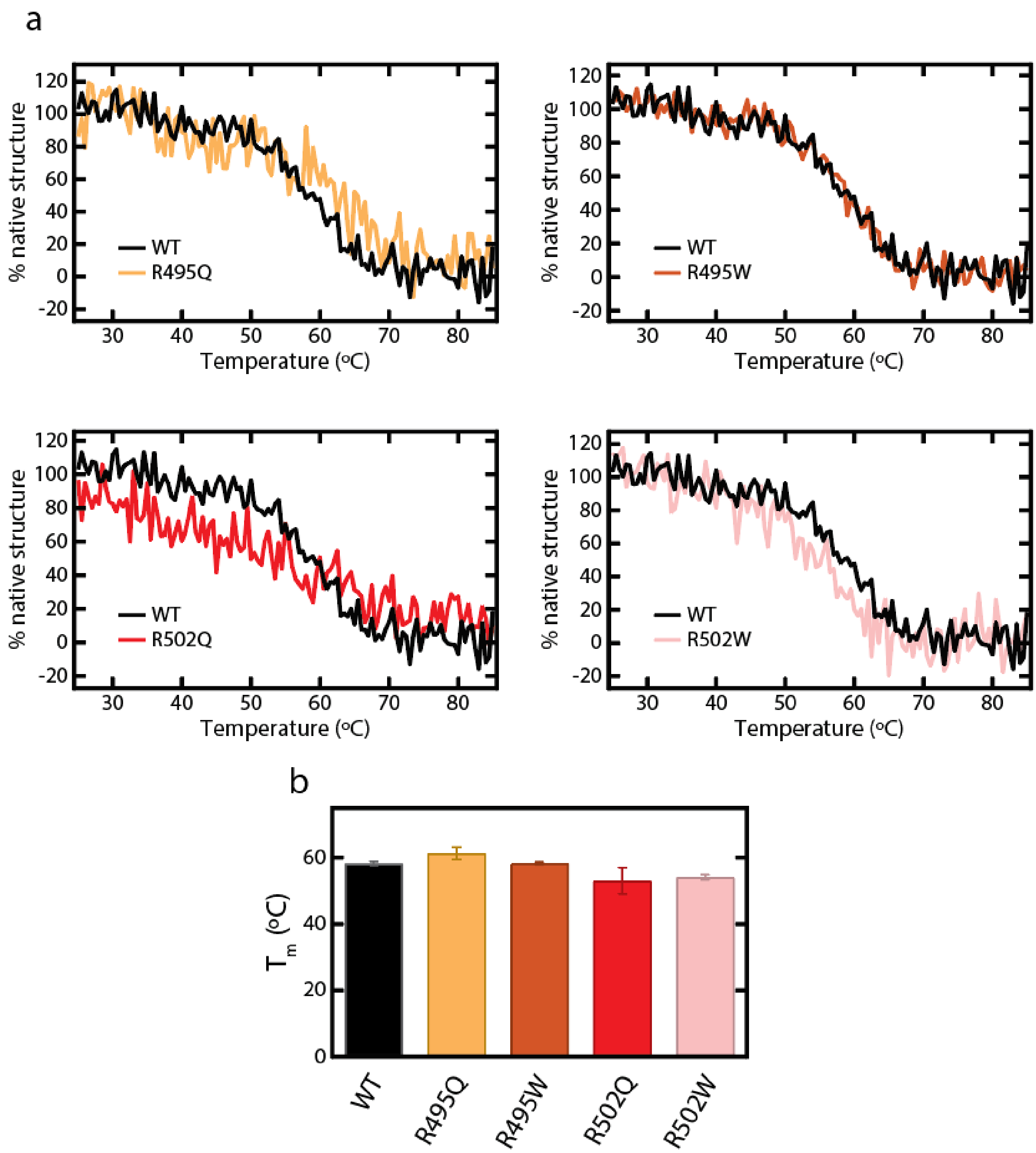


**Supplementary Figure S4. Thermal stability of WT and mutant C3 domains. (a)** Thermal denaturation curves of WT and mutant C3 domains obtained by tracking the CD signal at a wavelength of 215 nm. The temperature at the midpoint of the transition, $T_m$, can be obtained by performing a sigmoidal fitting to denaturation curves considering a two-state unfolding process. **(b)** $T_m$ for WT and mutant C3 domains. Error bars correspond to 83% confidence intervals (see also **Supplementary File S1**).

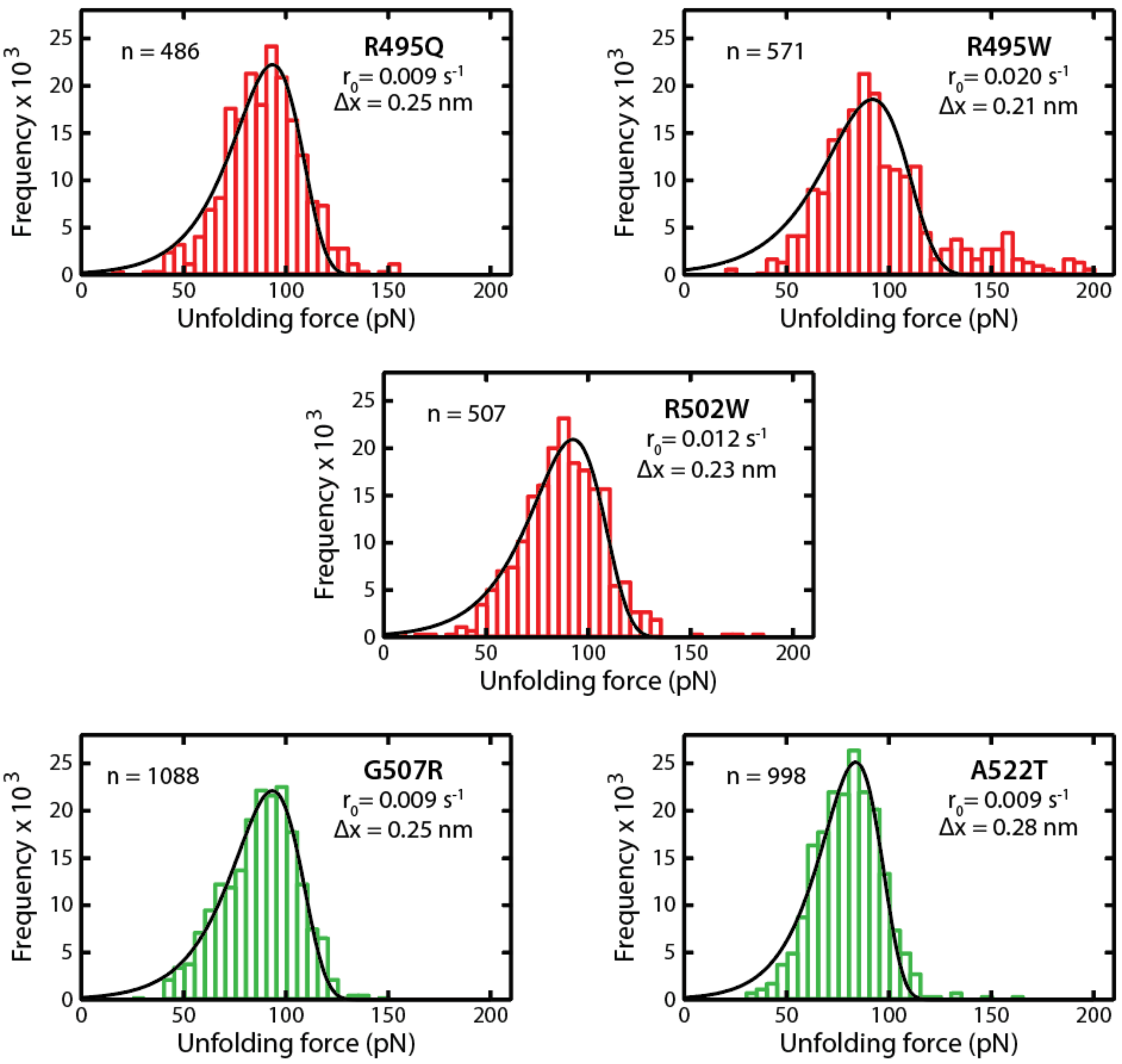


**Supplementary Figure S5. Distributions of unfolding forces for mutant C3 domains.** Distributions were fit to the Bell's model for force-activated protein unfolding [2]. Fitting lines appear in black and resulting parameters ($r_0$ and $\Delta x$) are shown. Distributions are colored according to the pathogenic status of the mutation (pathogenic, red; non-pathogenic, green).

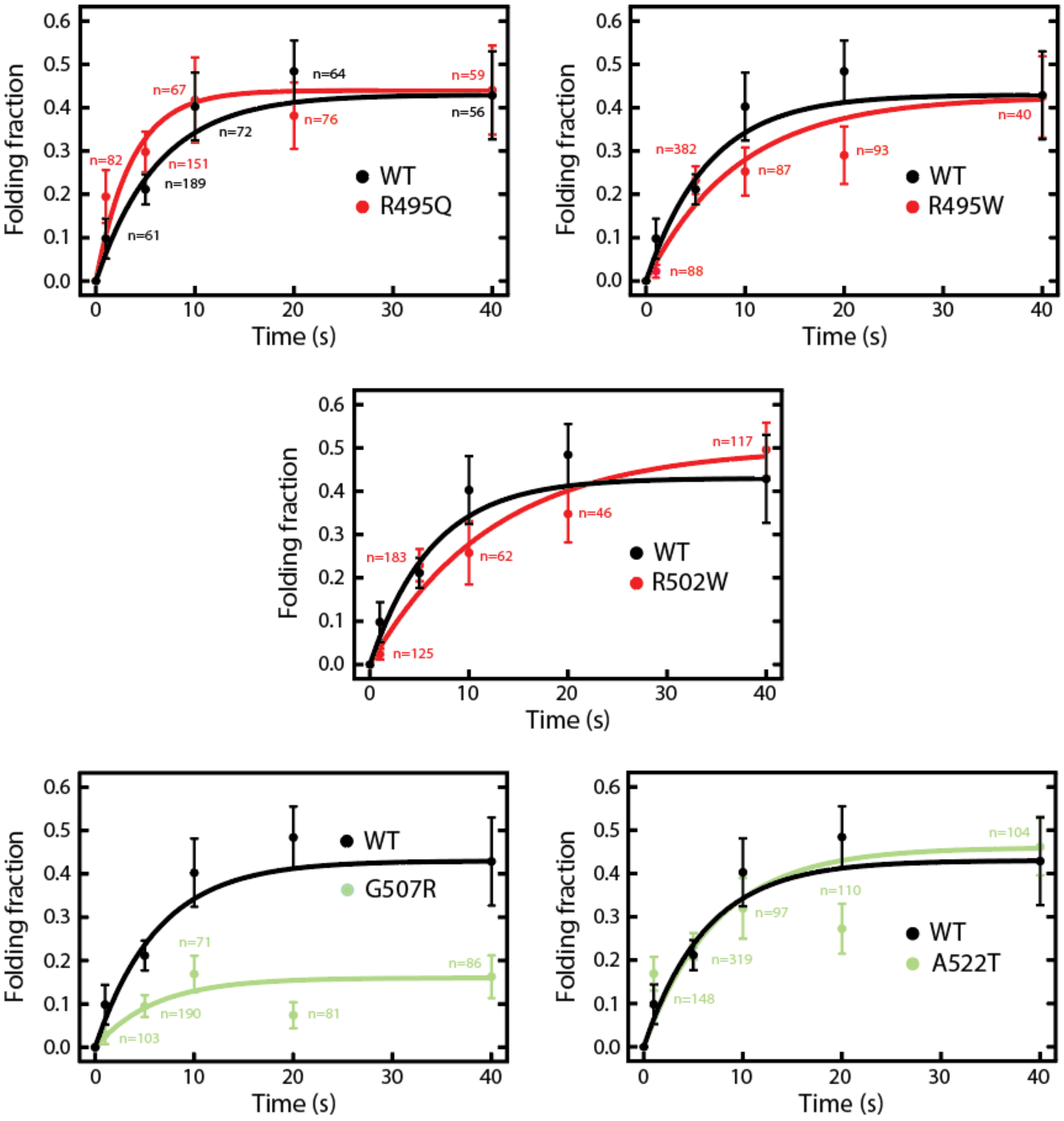


**Supplementary Figure S6. Folding kinetics.** Solid lines are exponential fits to the data. Error bars represent standard error of the mean, as estimated by bootstrapping. The number of unfolding events used to calculate folding fractions are indicated. Data points and fitting lines are colored according to the pathogenic status of the mutation (pathogenic, red; non-pathogenic, green).
